# Rice embryogenic trigger BABY BOOM1 promotes somatic embryogenesis by upregulation of auxin biosynthesis genes

**DOI:** 10.1101/2020.08.24.265025

**Authors:** Imtiyaz Khanday, Christian Santos-Medellín, Venkatesan Sundaresan

## Abstract

Somatic embryogenesis, a powerful tool for clonal propagation and for plant transformation, involves cellular reprogramming of differentiated somatic cells to acquire pluripotency. Somatic embryogenesis can be induced by treating explants with plant growth regulators. However, several plant species including agronomically important cereal crops remain recalcitrant to dedifferentiation and transformation except from embryonic tissues. Somatic embryogenesis can also be induced by ectopic expression of select embryonic factors, including in cereals by *BABY BOOM (BBM)* transcription factors. How *BBM* genes bypass the need for exogenous hormones is not well understood. Here, we investigated downstream targets during induction of somatic embryogenesis in rice by *OsBBM1* ((*Oryza sativa BABY BOOM1*). Transient induction of *OsBBM1* led to the upregulation of auxin biosynthesis *OsYUCCA* genes. Continued induction of *OsBBM1* resulted in somatic embryogenesis without the need for exogenous auxins. Genetic mutant analysis of *OsBBM1* downstream targets, *OsYUCCA6, OsYUCCA7* and *OsYUCCA9*, show that they are required for normal rice development including root and shoot development. Somatic embryogenic potential of *OsYUCCA* triple mutants was highly compromised despite the presence of exogenous auxin. Additionally, we show that somatic embryogenesis induction by exogenous auxin in rice requires functional *BBM* genes. Thus, *OsBBM1* mediated cellular reprogramming and somatic embryogenesis likely involves increased localized auxin through direct upregulation of *OsYUCCA* genes. This study reveals mechanistic details of how somatic embryogenesis is established in differentiated tissues in rice, a monocot model and agronomically important cereal crop, with the potential utility to improve regeneration from tissue culture for recalcitrant plants in future.

**One-sentence summary:** Rice BABY BOOM1 induces somatic embryogenesis from differentiated tissues by promoting auxin biosynthesis through direct upregulation of *YUCCA* genes.

## INTRODUCTION

Plant cells possess the remarkable property of dedifferentiation, the ability to return to a meristematic state whereby they can be redirected to various developmental fates including pluripotency and totipotency. This can be achieved by endogenous or exogenous stimuli such as the expression of various pluripotency factors or by treatment with plant hormones and stresses (Elhiti et al., 2013; Su et al., 2020). Somatic embryogenesis is a method of clonal propagation and a powerful tool for dedifferentiation and subsequently plant regeneration. Although different ways of inducing somatic embryogenesis exist in dicots (Mendez-Hernandez et al., 2019), many plant species including monocots and especially agronomically important cereal crops remain resistant to *in vitro* tissues culture, except for embryonic tissues (Eudes et al., 2003). A number of plant genes such as *BABY BOOM* (*BBM*) (Boutilier et al., 2002), *LEAFY COTYLEDON1* (*LEC1*), *LEC2* (Lotan et al., 1998; Stone et al., 2001; Wojcikowska et al., 2013), *WUSCHEL* (*WUS*) (Zuo et al., 2002), *FUSCA3* (*FUS3*) (Luerssen et al., 1998), *ABA INSENSITIVE3* (*ABI3*) (Parcy et al., 1994; Shiota et al., 1998), *EMBRYOMAKER* (Tsuwamoto et al., 2010)*, AtMYB115, AtMYB118* (Wang et al., 2009)*, SOMATIC EMBRYOGENESIS RECEPTOR*-*LIKE KINASE* (*SERK*) (Schmidt et al., 1997) and *AGAMOUS*-*LIKE 15* (*AGL15*) (Harding et al., 2003) have been shown to either induce or promote somatic embryogenesis in dicotyledonous plants. Among plant hormones, treatment with auxin is most widely used to induce totipotency and embryogenesis from differentiated plant cells (Wojcik et al., 2020). Stress conditions like osmotic stress, temperature stress, heavy metal stress, ultraviolet irradiation, wounding and chemical treatments have been used to promote somatic embryogenesis (Feher, 2015; Su et al., 2020).

*BABY BOOM* genes are members of the AINTEGUMENTA (ANT) clade of the superfamily of APETALA 2/ETHYLENE RESPONSE FACTOR (AP2/ERF) transcription factors with two AP2 domains (Dipp-Alvarez and Cruz-Ramirez, 2019). The first member of this family was identified in *Brassica napus* microspore cultures and its ectopic expression induced somatic embryogenesis on seedlings (Boutilier et al., 2002). The property of *BBM* genes to induce somatic embryogenesis is well documented in several other dicot species (Jha and Kumar, 2018). While only one *BBM* gene has been identified in Arabidopsis, four *BBM*-like genes exist in rice (Dipp-Alvarez and Cruz-Ramirez, 2019; Khanday et al., 2019). In Arabidopsis, *BBM* has been shown to express in zygotic embryos (Boutilier et al., 2002; Galinha et al., 2007); however, the role of *AtBBM* in zygotic embryogenesis has not yet been characterized, and loss-of-function mutants show no phenotypic abnormalities (Galinha et al., 2007)). We previously showed that *BBM* genes play an important role in zygotic embryogenesis in rice; specifically, that paternal expression of BBM factors is essential for embryo initiation and three *BBM* genes, *OsBBM1*, *OsBBM2* and *OsBBM3* redundantly regulate embryo development (Khanday et al., 2019).

*YUCCA* (*YUC*) genes encode flavin-containing monooxygenase enzymes that catalyze the rate-limiting step of oxidative decarboxylation of indole-3-pyruvate acid during auxin biosynthesis to form indole-3-acetic acid (IAA), the predominant auxin found in plants (Cao et al., 2019). The Arabidopsis genome encodes 11 *YUCCA* genes, whereas 14 members of this family have been identified in rice (Gallavotti et al., 2008). *YUC* gene mediated auxin biosynthesis has been shown to play diverse roles during various phases of plant development and response to environmental changes (Cao et al., 2019). *YUC* genes have been shown to regulate root and shoot development (Chen et al., 2014; Zhang et al., 2018), leaf morphogenesis (Cheng et al., 2006, 2007), inflorescence development (Woodward et al., 2005; Gallavotti et al., 2008), floral organ development (Cheng et al., 2006), establishment of the embryonic axis (Robert et al., 2013) and embryo organ initiation (Cheng et al., 2007).

We previously reported that ectopic expression of *OsBBM1* can induce somatic embryogenesis in differentiated tissues in rice (Khanday et al., 2019), thus making it a potential model for inducing and understanding somatic embryogenesis in monocots. Many downstream target genes regulated by *At*BBM during somatic embryogenesis have been identified in Arabidopsis (Passarinho et al., 2008; Horstman et al., 2017). However, a specific mechanism by which BBM factors induce somatic embryogenesis and support a hormone-independent growth of explants remains to be fully characterized. More generally, the mechanisms underlying induction of somatic embryogenesis in monocots are not known. In this study, we characterized the downstream target genes and pathways regulated by rice BBM1. We show that *Os*BBM1 directly activates the expression of auxin biosynthesis *YUCCA* genes to promote somatic embryogenesis. We further show the *OsYUCCA* genes targeted by *Os*BBM1 are essential for plant development and somatic embryogenesis.

## RESULTS

### Global Profile of Genes Deregulated on Induction of *Os*BBM1

In order to understand the gene regulatory network underlying initiation of embryogenesis in somatic tissues by *BBM* genes, we translationally fused *OsBBM1* gene product to a dexamethasone (DEX) inducible rat glucocorticoid receptor (GR) (Supplemental Fig. S1A; Khanday et al., 2019). Of the 10 transgenic lines raised with this construct, line #5 was found to have the best expression for the transgene *GR* (Supplemental Fig. S1C), thus all the subsequent experiments were performed in the progenies from line #5. For controlled induction of *Os*BBM1 expression, we took two-week old seedlings, wild-type or transgenic for the *OsBBM1-GR* fusion and treated them for various time points with DEX or a protein synthesis inhibitor, cycloheximide (CYC) or a combination of both (Supplemental Fig. S1D; see Materials and Methods). We choose 6 h time point for treatments based on the induction of *OsYUC7*, a potential target gene of *Os*BBM1 (Supplemental Fig. S1B and S1D). *OsYUC7* expression was induced in rice zygotes soon after fertilization and this expression pattern coincides with expression of *OsBBM1* (Anderson et al., 2017). *OsYUC7* expression was also found to be highly induced in *OsBBM1* overexpression lines in rice (Supplemental Fig. S1B; Khanday et al., 2019). Among the various time points tested, *OsYUC7* attained peak induction in 6 h time point, thus this time point was chosen for gene expression analysis regulated by *Os*BBM1 (Supplemental Fig. S 1D).

From our RNA-seq analysis of the samples treated with DEX for 6 hours, we detected a total of 298 differentially expressed (DE) genes in *OsBBM1-GR* seedlings at a log_2_ fold change of >1 and FDR < 0.05, after normalization with mock and wild-type controls (see Materials and Methods) (Supplemental Data Set S1). Of these 298 DE transcripts, 162 were upregulated and 136 were downregulated on *Os*BBM1 induction (Fig. 1, A and B). In the upregulated category, the expression of 138 genes was maintained in the presence of CYC, suggesting that they might be direct targets of *Os*BBM1 (Fig. 1A; Supplemental Data Set S2). From the downregulated category, the expression of 120 transcripts was also suppressed in the presence of CYC, which indicates *Os*BBM1 can directly represses the expression of these genes (Fig. 1B; Supplemental Data Set S2). A closer inspection of the genes directly regulated by *Os*BBM1 shows they belong to different functional categories including transcription factors, cell signaling, cell cycle regulation, hormone signaling, metabolism etc. (Supplemental Data Set S1 and S2). We confirmed the perturbations in expression of some the representative members of these different functional categories of genes in both upregulated and downregulated genes by RT-qPCR (Fig. 1, C and D).

**Figure 1.**
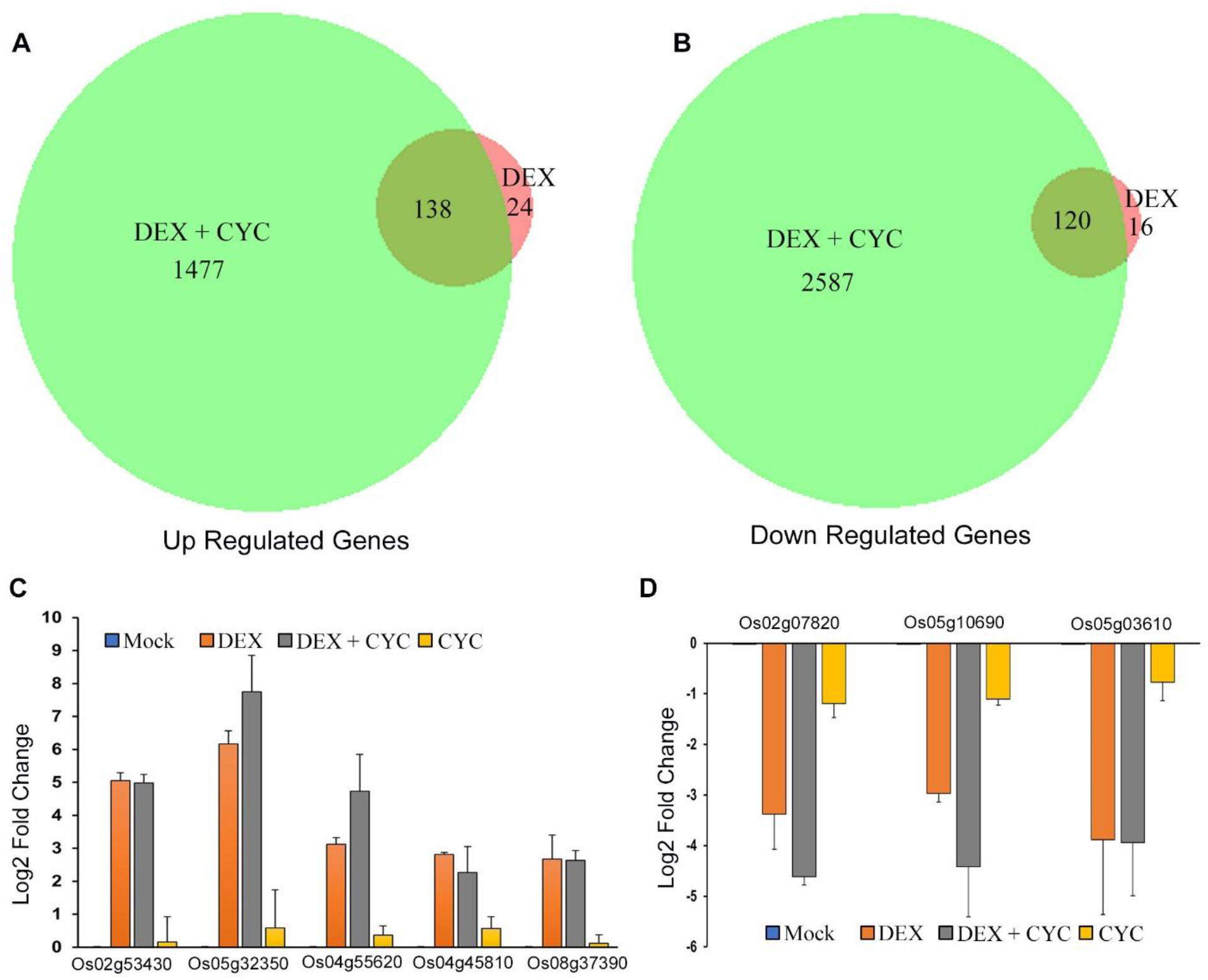
Changes in global gene expression in rice plants on *Os*BBM1-GR induction. A, RNA-sequencing analysis showing number of genes up regulated on induction of *Os*BBM1-GR with DEX alone or in the presence of CYC. B, A Venn diagram showing the down regulated transcripts by *Os*BBM1-GR in rice leaves. The numbers are genes deregulated by *Os*BBM1-GR under each treatment (Log_2_ fold >1 and FDR < 0.05). C and D, RT-qPCR validation of some of the genes regulated by *Os*BBM1. C, Genes from up regulated category are *Os04g45810*, a homeobox leucine zipper transcription factor, *Os05g32350*, a zinc finger containing protein, *Os04g55620*, a receptor kinase, *Os08g37390*, a cyclin and *Os02g53430*, a DNA-3-methyladenine glycosylase. D, From the down regulated category, the genes are *Os05g10690* is a MYB domain transcription factor, *Os02g07820* a DUF581 domain containing protein and *Os05g03610* is phospholipase C protein. The log_2_ fold change is calculated after normalization with mock and respective wild-type treatments. Error bars represent SEM, calculated from two independent biological replicates, each done in triplicates. DEX and CYC treatments were done for 6 h (see Materials and Methods). DEX, dexamethasone and CYC, cycloheximide.

### *Os*BBM1 Upregulates Expression of Auxin Biosynthesis Genes

Plant hormone auxin is known to regulate various developmental and physiological processes (Cao et al., 2019). Among the genes whose expression is regulated by *Os*BBM1, we find members of the *YUCCA* family of auxin biosynthesis genes and some auxin response factors (Supplemental Data Set S1). We confirmed the upregulation of expression for three *OsYUCCA* genes, *OsYUC6* (*Os07g25540*), *OsYUC7* (*Os04g03980*) and *OsYUC9* (*Os01g16714*) by RT-qPCR (Fig. 2A). The elevation in the expression of all the three genes by DEX was also maintained in the presence of CYC, suggesting their direct regulation by *Os*BBM1 (Fig. 2A). To further confirm that changes in the expression of these auxin biosynthesis genes are directly modulated by *Os*BBM1, we probed the occupancy of *Os*BBM1 at *cis*-regulatory regions of these Os*YUC* genes. Chromatin immune-precipitation (ChIP) was carried out with an anti-GR antibody (Sorefan et al., 2009), with tissues from *OsBBM1-GR* DEX treated plants of similar age as used for the RNA-seq experiment (see Materials and Methods). ChIP PCR reactions were carried out for 2 kb upstream sequences using various primers combination (Supplementary Table S2). *Os*BBM1 occupancy was observed in a region about 1.4 kb upstream of the transcription start site (TSS) of *OsYUC6* (Fig. 2B), in the immediate upstream sequences of *OsYUC7* (Fig. 2C) and about 1.1 kb upstream of TSS for *OsYUC9* (Fig. 2D). Significant enrichment at these sites was further confirmed by ChIP-qPCR over wild-type and input controls (Fig. 2E). The other regions within these 2 kb upstream sequences showed no enrichment by ChIP-PCR (Supplemental Fig. S2, A - C). A closer inspection of the sequences of *Os*BBM1 binding sites in the *cis*-regulatory regions of these *OsYUCCA* genes shows they contain putative sequence (CACA (N)_5_CNA or reverse complement GTGT(N)_5_GNT) of *At*BBM binding site (Horstman et al., 2017). Two of these sequence elements are present in *Os*BBM1 bound regions of *OsYUC7* and *OsYUC9,* whereas only one site is present in *OsYUC9.*Such sequences are also present in the other upstream regions of these *OsYUCCA* genes; however, we did not observe any *Os*BBM1 occupancy on those sites (Supplemental Fig. S2, A - C). Thus*, Os*BBM1 enrichment is site specific in the *cis*-regulatory regions of these *OsYUCCA* genes. Together with *Os*BBM1-GR DEX-induction data, these results confirm that *Os*BBM1 directly regulates the expression of *OsYUC6, OsYUC7* and *OsYUC9* auxin biosynthesis genes.

**Figure 2.**
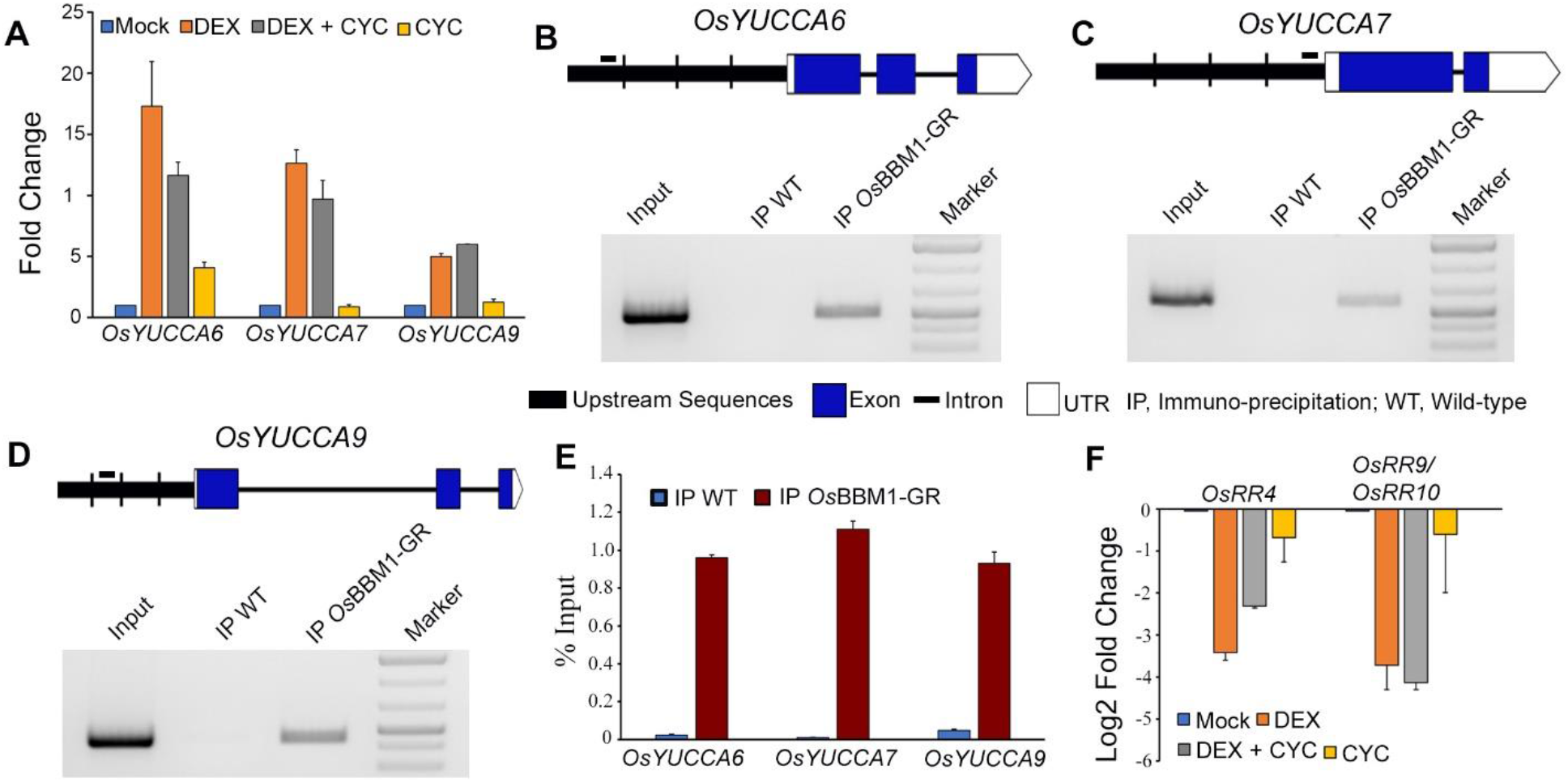
Regulation of plant hormone pathway genes by *Os*BBM1. A to E, Direct up regulation of auxin biosynthesis genes by *Os*BBM1. A, RT-qPCR analysis showing quantification of transcript levels for auxin biosynthesis genes *OsYUC6*, *OsYUC7* and *OsYUC9* in *OsBBM1-GR* plants. Fold change in normalized expression is shown for samples treated for 6 h with DEX, CYC or a combination of both the chemicals. The expression levels are normalized with mock treated samples and also with the similar treatments in wild-type tissues. Error bars were calculated as SEM, from two independent biological replicates. B to D, Chromatin-immunoprecipitation (ChIP) assay for enrichment of *Os*BBM1 at three *OsYUCCA* loci. ChIP-PCRs are shown for (B) *OsYUC6*, (C) *OsYUC7* and (D) *OsYUC9*. Schematics show the genomic organization of each locus. Exons are represented by blue box, introns by thin black line, UTRs with unfilled boxes and upstream sequences with thick black line. The short bar above the schematic shows the position of *Os*BBM1 occupancy on upstream sequences. E, ChIP-qPCR for quantitation of *Os*BBM1 enrichment on the three *OsYUCCA* loci. The enrichment is presented as percentage of input DNA. Error bars are represented as SEM, from two independent biological replicates. F, Quantitation of downregulation in expression of cytokinin response genes by RT-qPCR. Changes in expression are calculated as described for (A).

### *Os*BBM1 Expression Phenocopies Auxin Induced Somatic Embryogenesis

Auxin treatment induces somatic embryogenesis during plant regeneration (Wojcik et al., 2020). It is required for cellular reprogramming and induction of embryogenic callus from wild-type seeds in rice (Wani et al., 2011). Ectopic expression of *OsBBM1* resulted in somatic embryo formation on rice leaves in the absence of auxin (Khanday et al., 2019). This result, combined with our observation that *OsBBM1* can directly upregulate the expression of *OsYUC* genes, led us to examine if *Os*BBM1 promotes somatic embryogenesis by controlling auxin biosynthesis. We germinated *OsBBM1-GR* seeds on mock and DEX containing media. While mock treated seeds grew normally (Fig. 3A), we observed callus induction from the scutellum tissues in seeds grown on DEX containing media (Fig. 3, B and C). The seedlings from zygotic embryos in DEX treated seeds germinated normally; however, they were shorter and twisted compared to the mock treated seeds (Fig. 3, A and B; Supplemental Fig. S3B). Interestingly, wild-type seeds germinated on auxin-rich media, in addition to callus formation from scutellum, also exhibited the twisted seedling phenotype (Supplemental Fig. S3A). Callus formation was also observed from the first two juvenile leaves in the DEX treated *OsBBM1-GR* seedlings or from any leaf that was in contact with the DEX containing media (Fig. 3D, Supplemental Fig. S3, C and D). However, no such phenotype was observed in wild-type seeds germinated on auxin containing media (Supplemental Fig. S3A). When the *Os*BBM1-GR induced calli were allowed to grow on the DEX containing media for 3 to 4 weeks, they went on to start organogenesis ultimately regenerated roots and form green foci, indicative of start of shoot regeneration (Fig. 3, E and F). However, the greening calli ultimately reverted back to proliferation phase with loss of chlorophyll and never completed shoot regeneration. Also, regenerated roots or roots from the zygotic embryos after a prolonged time (3-4 weeks) on the DEX containing media started to dedifferentiate into callus-like tissues (Supplemental Fig. S3E). Thus, *OsBBM1* expression is sufficient to compensate for the auxin requirement for inducing somatic embryogenesis. In addition, when one-week old *OsBBM1-GR* seedlings grown on mock media were shifted to a DEX containing medium, they developed roots from aerial tissues (Supplemental Fig. S3, F and G). Moreover, *OsBBM1-GR* plants watered with DEX around the flowering transition developed awns from lemmas, which is not observed in wild-type rice plants (Supplemental Fig. S3, H and I). These vegetative and floral phenotypes have been shown to be associated with mutations related to auxin signaling and response (Toriba and Hirano, 2014; Mignolli et al., 2017).

**Figure 3.**
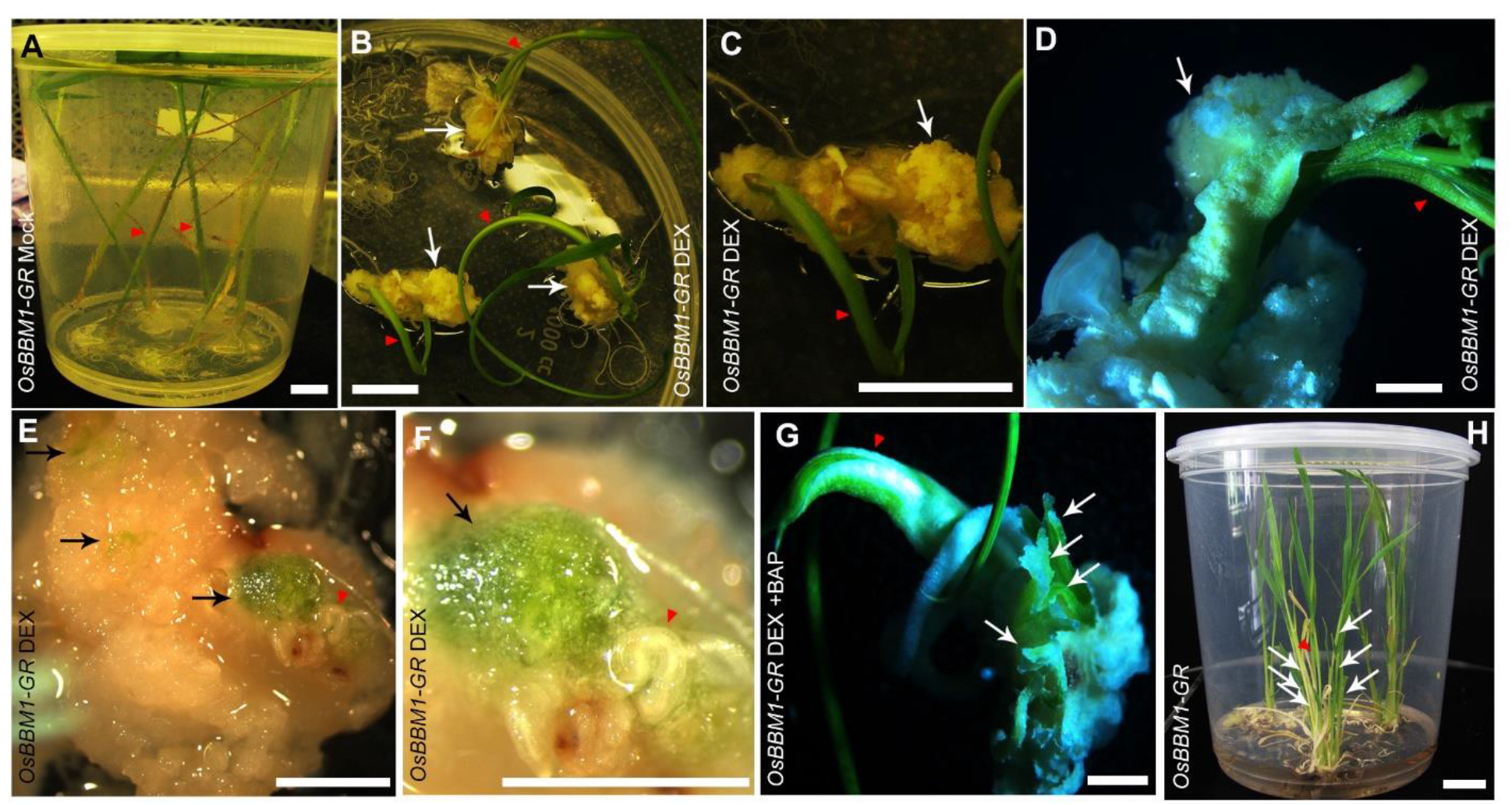
Phenotypic consequences of up regulation of auxin biosynthesis by *Os*BBM1 in rice seeds. A, *OsBBM1-GR* seeds germinated on mock treated medium. Seedlings grow normally (red arrowhead). B and C, *OsBBM1-GR* seeds grown on dexamethasone (DEX) containing media showing callus formation from scutellum tissue of seeds (white arrow). Red arrowhead points to the normally growing zygotic seedling. D, Somatic embryo formation from leaves of an *OsBBM1-GR* seedling induced with DEX (white arrow). Red arrowhead shows the growing zygotic seedling. E and F, Regenerating *OsBBM1-GR* calli induced with DEX. Black arrow shows the chlorophyll/shoot development and red arrowhead marks the regenerating roots. G, Complete plantlet formation from *Os*BBM1-GR DEX induced somatic embryos with addition of cytokinin. White arrows show the regenerating plantlets from somatic embryos and red arrowhead shows the seedling growing from the zygotic embryo. BAP is 6-Benzylaminopurine. H, Complete development of DEX+BAP regenerated plantlets in (G), when taken off both the treatments. Red arrowhead points to zygotic seedling and white arrows show regenerated plantlets from somatic embryos. Scale bars are 1 cm in (A), (B), (C) and (H), and 2 mm in (D to G).

We next investigated if externally applied auxin requires *Os*BBM1 for initiating somatic embryogenesis, by subjecting *Osbbm* mutant seeds to auxin treatment. Three rice *BBM* genes, *OsBBM1*, *OsBBM2* and *OsBBM3* have been shown to redundantly regulate zygotic embryo development (Khanday et al., 2019). We germinated seeds on callus induction medium from a mother plant segregating for *OsBBM1* (*OsBBM1*/*Osbbm1*) but homozygous mutant for *OsBBM2* (*Osbbm2*/*Osbbm2*) and *OsBBM3* (*Osbbm3*/*Osbbm3*) (see Materials and Methods). While all the control wild-type seeds (n=25) formed callus on auxin rich media (Supplemental Fig. S4A), some of the segregating mutant seeds failed to undergo any somatic embryogenesis (Supplemental Fig. S4B, red arrows). After genotyping (Supplemental Fig. S4F), seeds forming callus were found to be either wild-type (*OsBBM1*/*OsBBM1 Osbbm2*/*Osbbm2 Osbbm3*/*Osbbm3;* n=6/26) or heterozygous (*OsBBM1*/*Osbbm1 Osbbm2*/*Osbbm2 Osbbm3*/*Osbbm3;* n=13/26) for *OsBBM1* (Supplemental Fig. S4, B, C, E and F). None of the homozygous mutant seeds (*Osbbm1*/*Osbbm1 Osbbm2*/*Osbbm2 Osbbm3*/*Osbbm3;* n=7/26) underwent somatic embryogenesis (Supplemental Fig. S4, B, D, E and F). Thus, auxin mediated induction of somatic embryogenesis displays a requirement for functional *BBM* genes.

### Functional Characterization of *OsYUUCA6*,*OsYUCCA7* and *OsYUCCA9* Genes

Our results that *Os*BBM1 can directly regulate the expression of three *OsYUCCA* genes (Fig. 2, A-E) combined with auxin perturbation phenotypes observed on induction of *Os*BBM1 (Fig. 3, Supplemental Fig. S3), led us to investigate the functions of these *YUCCA* genes in rice and decipher their role in promoting embryogenesis downstream of *Os*BBM1. We created mutants in these three *OsYUCCA* genes using genome editing (see Materials and Methods). We created an *Osyuc7* Os*yuc9* double mutant and an *Osyuc6 Osyuc7* Os*yuc9* triple mutant (Supplemental Fig. S5 and S6). Ten independent transgenic lines with *CRISPR-Cas9* construct were raised for *Osyuc7* Os*yuc9* double mutant and 12 for *Osyuc6 Osyuc7* Os*yuc9* triple mutant. Two randomly selected lines were used for detailed mutation analysis for each mutant combination (Supplemental Fig. S5 and S6). T2 progenies in which *CRISPR-Cas9* transgene had already segregated out (Supplemental Fig. S5C) and were null mutations for all alleles of all the genes (Supplemental Fig. S5, A, B and S6, C and D) were taken further for phenotypic analysis.

Mutations in Os*YUC* genes affected various vegetative characteristics of rice plants. Plants double mutant for *OsYUC7* and *OsYUC9* were shorter compared to the wild-type plants (Fig. 4, A and E). The phenotype of reduction in height was further enhanced in *Osyuc6 Osyuc7* Os*yuc9* triple mutant plants (Fig. 4, B and E). Another prominent phenotype in these mutant plants was elongation in primary root length (Fig. 4, C and D). While average root length in 12 day old wild-type control seedlings was 4.4 cm, it increased to 15.3 cm in *Osyuc7* Os*yuc9* double mutant (Fig. 4, C and F). This phenotype was further elevated in *Osyuc6 Osyuc7* Os*yuc9* triple mutant (Fig. 4D). The average root length in *Osyuc6 Osyuc7* Os*yuc9* triple mutant plants was about 23.5 cm (Fig. 4F). Lateral root development was also impaired in rice *YUC* mutants. Compared to the wild-type, fewer lateral roots developed in *Osyuc7* Os*yuc9* double mutant (Supplemental Fig. S5, D and E). This lateral root development defect was more prominent in *Osyuc6 Osyuc7* Os*yuc9* triple mutant (Supplemental Fig. S5F).

**Figure 4.**
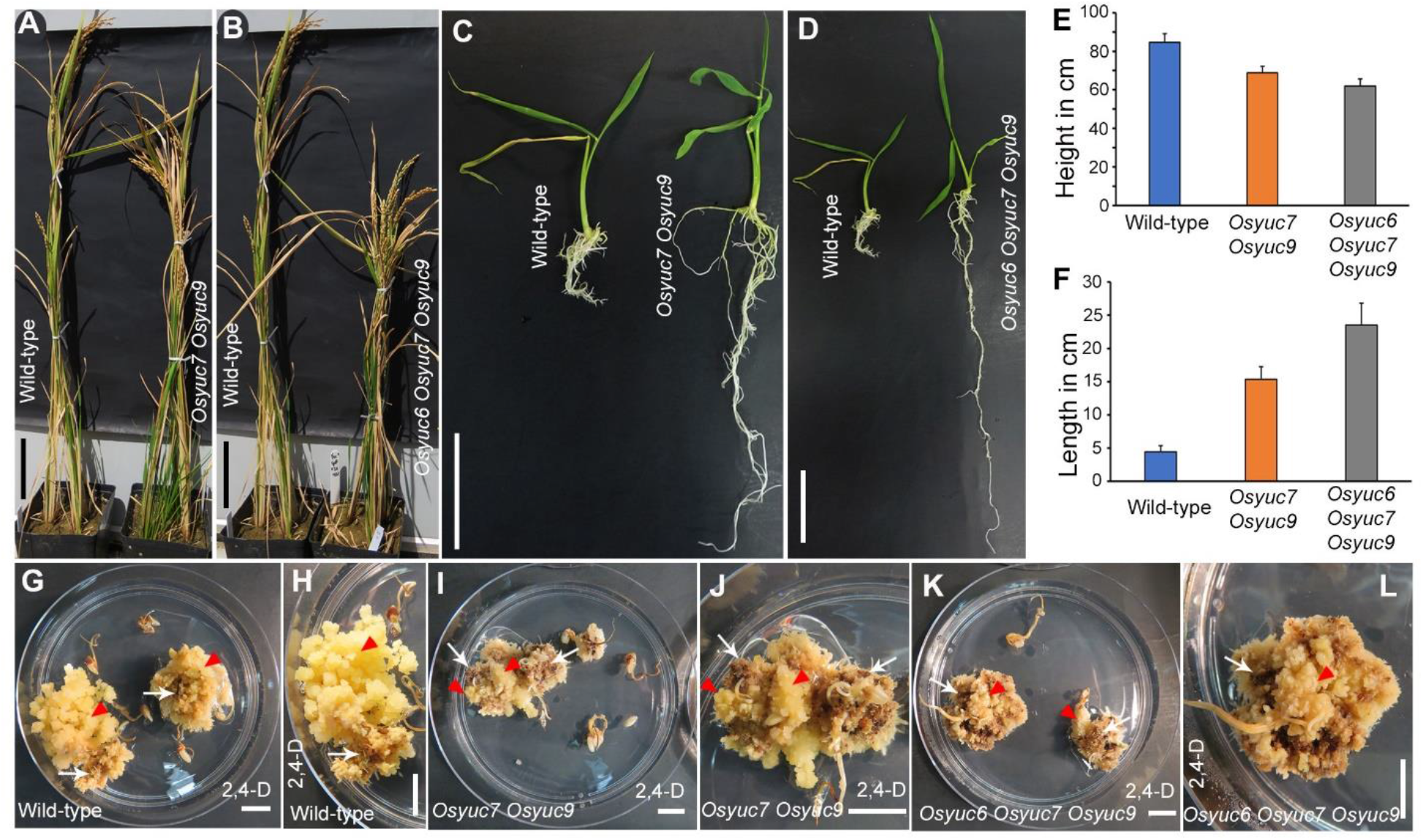
Functional characterization of *OsYUCCA6*, *OsYUCCA7* and *OsYUCCA9* genes in rice. A and B, Reduction in plant height in *Osyucca* mutants. A, *Osyucca7 Osyucca9* double mutant compared with wild-type. B, *Osyucca6 Osyucca7 Osyucca9* triple mutant compared to wild-type. Scale bars 10 cm. C and D, Perturbations in primary root length in *Osyucca* mutants. C, *Osyucca7 Osyucca9* double mutant. D, *Osyucca6 Osyucca7* Os*yucca9* triple mutant. Scale bars 5 cm. E, Bar graph showing the quantification of reduction in plant height in *Osyucca* mutants (n=20 for each genotype). F, Quantitative representation of increase in root length in *Osyucca* mutants in rice (n=20 for each genotype). G to L, Reduction in embryogenic calli induction in *Osyucca* mutants in the presence of supplemented auxin. G and H, Wild-type showing induction and proliferation of compact, pale and nodular embryogenic calli (red arrow heads) which are phenotypically distinct from brown sectors of non-embryogenic calli (white arrows). I and J, *Osyucca7 Osyucca9* double mutant showing some proliferation of embryogenic calli (red arrow heads). K and L, *Osyucca6 Osyucca7 Osyucca9* triple mutant, showing extensive brown sectors of non-embryogenic calli (white arrows) without proliferation of pale embryogenic calli (red arrowheads). 2,4-D is 2,4-Dichlorophenoxyacetic acid. Scale bars G to L are 1 cm.

Next, we analyzed the effect of these *OsYUCCA* mutations on embryogenic callus formation. Seeds from wild-type controls, *Osyuc7* Os*yuc9* double and *Osyuc6 Osyuc7* Os*yuc9* triple mutant plants were germinated in dark on media containing 2 mg/L of 2,4-Dichlorophenoxyacetic acid (2,4-D) for callus induction. While we observed robust induction and proliferation of calli from the embryonic tissues in wild-type seeds (n=37/60) (Fig. 4, G and H), embryogenic calli (nodular, compact and pale) formation was severely reduced in *Osyuc7* Os*yuc9* double mutant seeds (n=8/60) (Fig. 4, I and J). In *Osyuc6 Osyuc7* Os*yuc9* triple mutant seeds, embryogenic calli induction was drastically reduced and exhibited almost no proliferation (n=3/60) (Fig. 4, K and L). Thus, despite supplementing the media with auxin to complement the auxin biosynthesis deficiency caused by the *OsYUCCA* mutations, callus induction in these mutants was severely compromised.

### Cytokinin Supplementation Completes Shoot Regeneration in *Os*BBM1 Induced Somatic Embryos

Cytokinin plant growth regulators are required for the shoot regeneration in tissue culture (Hill and Schaller, 2013). The DEX induction of *Os*BBM1-GR resulted in somatic embryos which started organogenesis, but shoot regeneration was not completed (Fig 3, E and F). Therefore, we tested the possibility of promoting the shoot regeneration process to completion, by supplementation of the media with cytokinin. Seeds germinated either in light or dark on media containing both DEX and cytokinin 6-Benzylaminopurine (BAP), showed regeneration of shootlets from somatic embryos induced by *Os*BBM1 expression (Fig. 3G, Supplemental Fig. S3J). When these regenerating plantlets were taken off both the treatments and grown on normal media, they completed the full developmental program and formed normal seedlings (Fig. 3H). From our RNA-seq analysis of DEX treated 6h leaf samples, we also found that *Os*BBM1 negatively regulates the expression of some cytokinin signaling genes, such as cytokinin response regulators like *OsRR4* (*Os01g72330*) and *OsRR9* (*Os11g04720*)/*OsRR10* (*Os12g04500*) (Fig. 2F and Supplemental Data Set S1 and S2). This putative repressive function is consistent with previous results from Arabidopsis BBM, which has been shown to interact with TOPLESS-related (TPR) corepressors (Causier et al., 2012), suggesting that BBM factors can also act as transcriptional repressors. Thus, these results demonstrate that supplementation of cytokinin can override the incomplete shoot formation in *Os*BBM1-induced embryogenesis, which might be related to *Os*BBM1-mediated suppression of cytokinin response factors.

## DISCUSSION

Although somatic embryogenesis has been a problem of long-standing interest in plant biology, it is not known whether a common mechanism underlies somatic embryogenesis between monocot and dicot species, due in part to limited studies on successful induction of somatic embryogenesis in monocot species. Previous studies in Arabidopsis show *At*BBM regulates somatic embryogenesis by regulating the expression of so called *LAFL* genes-*LEC1*, *LEC2*, *FUS3* and *ABI3* (Horstman et al., 2017). Further, *At*LEC1 can regulate *AtYUC10* expression (Junker et al., 2012) and *AtLEC2* expression has been shown to be associated with expression of *AtYUC1*, *AtYUC4* and *AtYUC10* genes (Wojcikowska et al., 2013). Although *At*BBM can bind to *cis*-regulatory sequences of *AtYUC3* and *AtYUC8* genes (Horstman et al., 2017), no changes in their expression or those of other *YUC* genes have been reported upon induction of somatic embryogenesis by *At*BBM (Passarinho et al., 2008). Our data show that in rice, *Os*BBM1 directly upregulates the expression of *YUCCA* genes-*OsYUC6*, *OsYUC7* and *OsYUC9* upon induction of somatic embryos (Fig. 2). These results may indicate differences in the mechanism between dicots and monocots in the control of auxin biosynthesis for induction of somatic embryos by *BBM* genes. Additionally, our analysis shows no detectable expression of either *OsLEC1* (*OsLEC1A*, *Os02g49370*; *OsLEC1B*, *Os06g17480*) or *OsLEC2* (*Os04g58000*, *Os04g58010* and *Os01g68370*) related genes in our leaf samples of 6 h DEX treatment (Supplemental Data Set S1). However, *OsLEC1A* and *OsLEC1B* expression was observed in somatic embryos established subsequent to induction by *OsBBM1* (Khanday et al., 2019). We note that changes in *AtLEC* gene expression were not observed in *35S:AtBBM-GR* DEX treated seedlings prior to formation of somatic embryos (Passarinho et al., 2008), although the *LAFL* genes were found to be targets of *At*BBM in somatic embryos as discussed above (Horstman et al., 2017). Therefore, it is possible that *LEC* genes are not required for the initiation of somatic embryogenesis in differentiated cells by BBM factors but are important during the later stages of somatic embryogenesis, possibly for maintenance of the embryonic state. The observation that *Os*BBM1 modulates auxins by regulating the expression of *OsYUC* genes is also supported by the fact that transient induction of *Os*BBM1-GR during other stages of development such as seedlings and flowers also leads to auxin perturbation phenotypes (Supplemental Fig. S3, F - I). These observations together with the phenotypes observed on *BBM1* overexpression (Khanday et al., 2019), support the evidence that *YUCCA* genes are immediate downstream targets of *Os*BBM1. Of the three *OsYUCCA* genes upregulated by *Os*BBM1 (Fig. 2), *OsYUC7* expression is *de novo* induced in rice zygotes after fertilization and peaks at 9 hours after pollination which correlates with *OsBBM1* expression (Anderson et al., 2017). Auxin biosynthesis genes have also been shown to be upregulated in maize zygotes after fertilization (Chen et al., 2017). These data suggest that the initiation of zygotic embryogenesis by BBM transcription factors might also involve the biosynthesis of auxin by direct activation of *YUCCA* genes. The differences in the spectrum of *YUCCA* genes activated during zygotic embryogenesis compared to the direct targets we observed in this study might reflect different chromatin configurations and relative accessibility of these genes in the two very different cell types, *i.e.* egg cells and leaf cells. The extent to which the pathways for the initiation of somatic and zygotic embryogenesis by BBM factors is conserved remains to be elucidated.

Tissue culture for plant regeneration is a two-step process in which explants are initially cultured on an auxin-rich medium to produce callus. These calli are then cultivated on a cytokinin-rich medium to regenerate shoots (Hill and Schaller, 2013). Although *Os*BBM1 was able to induce somatic embryogenesis, and embryos proceeded to greening and roots development, the entire process of complete plantlet regeneration was never achieved without the supplementation of media with cytokinin (Fig. 3). The down regulation of some cytokinin response regulator genes by *Os*BBM1 (Fig. 2F) might partially account for the requirement of cytokinin to complete the shoot formation in *Os*BBM1 induced somatic embryos.

In Arabidopsis, it was found that efficiency of somatic embryogenesis induced by external auxin is reduced by mutants in two *YUC* genes (Wojcikowska et al., 2013). We show here that in rice, induction of somatic embryogenesis is highly compromised in *Osyuc7* Os*yuc9* double and *Osyuc6 Osyuc7* Os*yuc9* triple mutant combinations, despite the presence of exogenous auxins in the medium (Fig. 4, I to L). The observation that endogenous IAA biosynthesis is required for somatic embryogenesis in rice and cannot be substituted by application of exogenous auxins is also consistent with the different developmental roles of endogenous and external auxin, deduced from previous studies in Arabidopsis. Root developmental defects caused by *yuc* mutations were not rescued by external application of auxins in Arabidopsis (Cheng et al., 2006; Chen et al., 2014). Thus, tissue-specific local auxin biosynthesis by *YUCCA* flavin monooxygenases is important both for plant development and somatic embryogenesis.

Auxin has been shown to be main inducer of somatic embryogenesis in various plant species (Wojcik et al., 2020). Auxin has also been shown to induce members of PLETHORA/BBM transcription factors in Arabidopsis roots (Mahonen et al., 2014), and also expression of *BBM* homologs in seedlings of the monocot cereal *Brachypodium distachyon* (Kakei et al., 2015). Differentiated leaves from rice do not express *OsBBM1* (Khanday et al., 2019) and are recalcitrant to callus induction even in the presence of auxin (Supplemental Fig. S3A; (Hu et al., 2017). However, rice leaves with ectopic expression of *OsBBM1,* or with *Os*BBM1-GR DEX-induction, undergo somatic embryogenesis (Fig. 3D; Khanday et al., 2019). These observations combined with our results that *Osbbm1 Osbbm2 Osbbm3* mutant seeds are unable to undergo somatic embryogenesis (Supplemental Fig. S4), suggest that externally applied auxin induces somatic embryogenesis by upregulating *BBM* gene expression in scutellar cells, but is unable to upregulate *BBM* gene expression in differentiated leaf cells (Fig. 5). Consistent with these observations, regeneration from tissue culture of a triple knock-out mutant of *OsBBM1*, *OsBBM2* and *OsBBM3* was never achieved, even though *OsBBM1-3* genes are dispensable for vegetative growth and development (Khanday et al., 2019).

**Figure 5.**
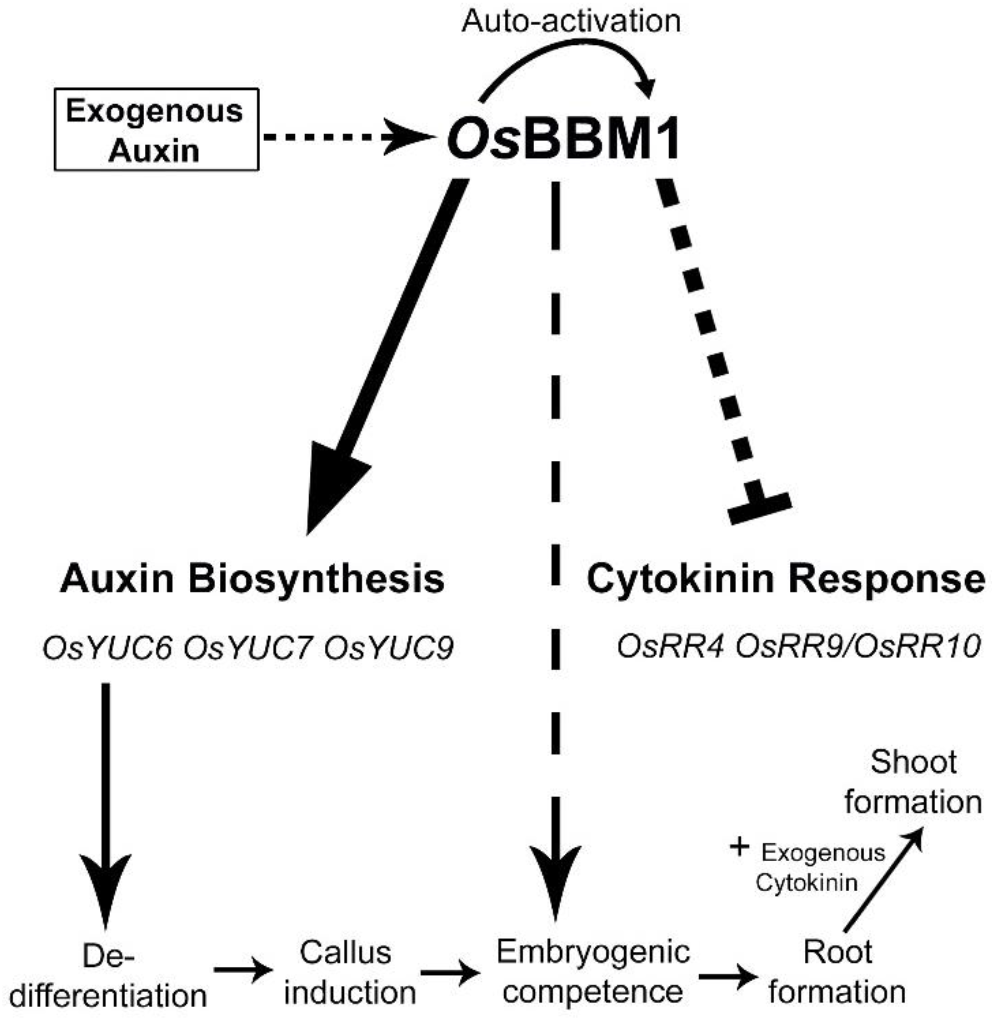
Model explaining the mechanism of *Os*BBM1 action during somatic embryogenesis in rice. *Os*BBM1 expression is required for dedifferentiation of somatic cells. In scutellum cells, *Os*BBM1 is downstream of exogenous auxin (dotted arrow), but in differentiated leaf cells that are recalcitrant to external auxin, *Os*BBM1 has to be expressed using a transgene as in this study. *Os*BBM1 positively regulates the expression of auxin biosynthesis *OsYUCCA* (*OsYUC6*, *OsYUC7* and *OsYUC9*) genes. This leads to increase in the endogenous auxin levels, followed by dedifferentiation, callus formation and subsequently somatic embryogenesis. *Os*BBM1 also auto-activates its promoter to maintain expression and pluripotency during somatic embryogenesis. The somatic embryos are able form roots and initiate shoot regeneration, but shoot formation is not completed, possibly due to negative regulation of some cytokinin response genes by *Os*BBM1. This block can be overcome by the addition of cytokinin in the culture media and complete plant regeneration is achieved.

In conclusion, *Os*BBM1 expression in differentiated tissues induces somatic embryogenesis in the absence of exogenous auxins by promoting local auxin biosynthesis through control of *OsYUCCA* gene expression. This function of *Os*BBM1 suggests a pathway for induction of somatic embryogenesis that may be applicable to cereals generally, as summarized in Figure 5. In scutellum or other tissues that are competent for somatic embryogenesis induction by exogenous auxin, auxin induces expression of *OsBBM1* (and presumptively, the closely related paralogs *OsBBM2* and *OsBBM3*). Loss of these *BBM* genes results in failure of external auxin to induce somatic embryos. In differentiated leaf, a tissue that is recalcitrant to somatic embryo induction by external auxin, the ectopic expression of *OsBBM1* is sufficient to activate the downstream somatic embryogenesis pathway. In both cases, *OsBBM1* expression, promotes endogenous auxin biosynthesis through *YUCCA* genes, as well as auto-activation of its own promoter to sustain its expression during somatic embryo formation (Khanday et al., 2019). External auxin cannot substitute for the crucial role of the *Os*BBM1-targeted *YUCCA* genes, as shown by mutant analysis. Elevated intracellular levels of IAA from the *YUCCA* genes leads to dedifferentiation, and somatic embryos that produce roots and greening callus with incipient shoot formation. However, complete formation of shoots requires exogenous cytokinin, following which viable plants can be regenerated. This initial characterization of the molecular pathway for induction of somatic embryogenesis in a cereal crop can help pave the way for future experiments to develop methods for efficient tissue culture and transformation in cereals and other recalcitrant plants.

## MATERIALS AND METHODS

### Plant Materials and Growth Conditions

All the experiments in this study were performed in rice cultivar Kitaake (*Oryza sativa* L. subsp. *japonica*). All the seeds, wild-type, transgenic and mutant were initially germinated on half strength Murashige and Skoog’s medium (Murashige and Skoog, 1962) containing 1% sucrose and 0.3% phytagel. After 12-15 days of initial growth on media, seedlings were transferred to greenhouse and grown there until seeds matured.

### Chemical Treatments

Dexamethasone (DEX) and cycloheximide (CYC) treatments were carried in two-week-old wild-type and *OsBBM1:GR* seedlings as described in Khanday et al., 2019. Leaf tissue samples for RNA isolation for each biological replicate were collected from 4 plants and pooled together. Tissues samples were harvested at 3h, 6h, 12h and 24 h after the start of treatment. For DEX induced phenotypes (Fig. 3, A-F and Supplemental Fig. S3, B-E) *OsBBM1-GR* seeds were grown under light on a media containing either mock or 10 μm DEX. In Fig. 3G and Supplemental Fig. S3J, in addition to DEX, media also contained cytokinin 6-Benzylaminopurine at 3 mg/L concentration and seeds were grown either in light (Fig. 3G) or in dark (Supplemental Fig. S3J). For phenotypes in Supplemental Fig. S3, F and G, one-week-old *OsBBM1-GR* seedlings were transferred to half strength MS media either mock treated or containing 10 μM DEX and grown for one more week. For phenotypes observed in Supplemental Fig. 3, H and I, plants started receiving either mock or 10 μM DEX water around flowering transition (45 days after germination) until the panicles completely emerged. For phenotypes shown in Fig. 4, G to L, Supplemental Fig. S3A and Supplemental Fig. 4, wild-type, *Osbbm* or *Osyuc* mutant seeds were germinated on media containing 2 mg/L of 2,4-Dichlorophenoxyacetic acid (2,4-D). The seeds were allowed to grow for two weeks under light (Supplemental Fig. S3A), two weeks in the dark (Supplemental Fig. 4) and for six weeks in the dark (Fig. 4, G to L).

### Microscopy and Image Capture

Microscopic images were captured using a Zeiss Stemi SV11 stereo microscope fitted AxioCam HRC camera (Zeiss). AxioVision version 4.8 software (Zeiss) was used for image capture and processing. Images for plant phenotypes were taken with Cannon PowerShot SX60 HS digital camera. All the images and other data panels were assembled into article figures using Adobe Photoshop (Adobe Inc.).

### RNA-Seq Library Preparation and Sequencing

RNA isolation, quality assessment, quantification and library preparations were done as described (Anderson et al., 2017). Libraries were prepared from two biological replicates for each sample with 80 ng of input RNA, using NuGEN Ovation RNA-seq Systems 1-16 following manufacturer instructions. Samples were multiplexed and 8 libraries per lane were run on Illumina HiSeq platforms at UC Davis, Genome Center.

### RNA-Seq Analysis

Cutadapt (Martin, 2011) was used to remove 3’ adapters and quality-trim reads at a Phred quality threshold of 13. High-quality reads were then mapped to the *Oryza sativa* Nipponbare reference genome (MSU v7.0)(Kawahara et al., 2013) using Tophat2 (Kim et al., 2013) with the minimum and maximum intron sizes set at 20 and 15000 bases and the microexon search switched on. Mapped reads were assigned to the MSU v7.0 gene models (Phytozome release 323) using HTSeq. Differential expression analysis was performed using the edgeR package (Robinson et al., 2010). Effective library sizes were calculated using the trimmed mean of M-values (TMM) method and a quasi-likelihood (QL) negative binomial generalized linear model was fitted to the data (Chen et al., 2016). The contrasts of DEX, DEX + CYC and CYC with their respective mock samples were created for both wild-type and *OsBBM1-GR* genotypes. For each contrast, a QL F-test was performed to detect differentially expressed genes (FDR < 0.05). Genes induced by DEX or CYC treatments in wild-type were subtracted from the respective treatments in *OsBBM1-GR* samples.

### RT-PCR and RT-qPCR

All the cDNA synthesis, RT-PCRs, RT-qPCRs and analysis of RT-qPCRs were performed as described (Khanday et al., 2019). All the RT-qPCR amplifications were done in two biological replicates and each biological replicate was repeated as three technical replicates. All the primers used for RT-PCRs and RT-qPCRs are listed in Supplemental Table S1.

### Chromatin Immunoprecipitation (ChIP)

Chromatin Immunoprecipitation (ChIP) was performed as described (Khanday et al., 2013) with following modifications. Wild-type and *OsBBM1-GR* two-week-old rice seedlings were grown in 0.5X Mureshige and Skoog’s medium and treated with 10 μM DEX for 6 h. Leaf tissues were harvested, and cross linked with 1% formaldehyde. For each biological replicate, chromatin was prepared from 300 mg of tissues. Chromatin was sheared to ~500 bp size using a Covaris E220 instrument in 1 ml Covaris milliTUBE. The instrument settings for chromatin shearing were 140 peak incident power (PIP), 5% duty factor, 200 cycles per burst (CBP). Chromatic complexes were immunoprecipitated with 2 μg of anti-GR antibody (AB3580, Abcam) (Sorefan et al., 2009). Chromatin-antibody complexes were recovered using Protein A conjugated Dynabeads (Invitrogen, 10001D). Primers used for ChIP-PCRs and ChIP-qPCRs are listed in Supplemental Table S2.

### Vector Construction, *Osyucca* and *Osbbm* Mutant Generation, and Genotyping

The construction of *pZmUBI1*::*OsBBM1-GR* (Supplemental Fig. S1A) is described in Khanday et al., 2019. For generating *Osyuc7 Osyuc9* double and *Osyuc6 Osyuc7 Osyuc9* triple mutants, single guide RNAs (sgRNAs) for gene editing for *OsYUC6* (*Os07g25540*), *OsYUC7* (*Os04g03980*) and *OsYUC9* (*Os01g16714*) were designed using the webtool https://www.genome.arizona.edu/crispr/ as described (Xie et al., 2014). The sgRNA sequences are listed in Supplemental Table S3. Assembly of plasmid constructs for CRISPR-Cas9 editing with these sgRNAs and rice transformations were done as described (Khanday et al., 2019). Ten and twelve independent T0 transformants were generated for *pCRISPR*-*OsYUC7* + *OsYUC9* and *pCRISPR-OsYUC6* + *OsYUC7*+ *OsYUC9* constructs, respectively. These T0 plants were screened for mutations by genotyping PCRs, followed by sequencing of the amplicons. The primers for genotyping PCRs are listed in Supplemental Table S3. T2 progenies form two lines for each of the two *CRSIPR-Cas9* constructs were studied in detail. Only those progenies in which *CRISPR-Cas9* construct was no longer present (Supplemental Fig. S5C) and had all the requisite null alleles for mutations were considered for analysis (Supplemental Fig. S5 and S6). *Osbbm* mutant generation and genotyping are described in detail in Khanday et al., 2019. Briefly, a single bp deletion mutation in *OsBBM1* CDS disrupts a *SphI* site. The DNA fragment at the mutation site is amplified with the genotyping primers listed in Supplemental Table S3 and restriction digested with *SphI (*Supplemental Fig. S4F).

#### Accession Numbers

Accession numbers for the genes characterized in this work are listed in the main text. RNA sequencing data from this article will be uploaded to GenBank data library and accession number XXX updated after review.

## Supplemental Data

**Supplemental Figure S1.** DEX inducible gene expression system for *Os*BBM1.

**Supplemental Figure S2.** Occupancy of *Os*BBM1 on *YUCCA* gene loci.

**Supplemental Figure S3.** DEX induced phenotypes in *OsBBM1-GR* plants.

**Supplemental Figure S4.** Somatic embryogenesis in *Osbbm* mutants.

**Supplemental Figure S5.** Mutations in *Osyucca7 Osyucca9* double mutant plants.

**Supplemental Figure S6.** Mutations in *Osyucca6 Osyucca7 Osyucca9* triple mutants.

**Supplemental Data Set S1.** Genes deregulated by *Os*BBM1-GR in Mock vs. DEX treatment.

**Supplemental Data Set S2.** Genes directly regulated by *Os*BBM1-GR in Mock vs. DEX+CYC treatment.

**Supplemental Table S1.** Primers used for RT-PCRs and RT-qPCRs.

**Supplemental Table S2.** Primers used for ChIP-PCRs and ChIP-qPCRS.

**Supplemental Table S3.** Sequences of sgRNAs and genotyping primers.

## ACKNOWLEDGMENTS

We thank Alina Yalda for technical assistance with genotyping and green house transplantations.

## Notes

**Funding information**: C.S-M. acknowledges support from the University of California Institute for Mexico (UCMEXUS), Consejo Nacional de Ciencia y Tecnología (CONACYT), and Secretaría de Educación Pública (México). This research was funded by grants from the National Science Foundation (IOS-1547760), Innovative Genomics Institute and the U.S. Department of Agriculture (USDA) Agricultural Experiment Station (CA-D-XXX-6973-H).

